# Mu-opioids suppress GABAergic synaptic transmission onto orbitofrontal cortex pyramidal neurons with subregional selectivity

**DOI:** 10.1101/2020.05.12.091678

**Authors:** Benjamin K. Lau, Brittany P. Ambrose, Catherine S. Thomas, Min Qiao, Stephanie L. Borgland

## Abstract

The orbitofrontal cortex (OFC) plays a critical role in evaluating outcomes in a changing environment. Administering opioids to the OFC can alter the hedonic reaction to food rewards and increase their consumption in a subregion specific manner. However, it is unknown how mu-opioid signalling influences synaptic transmission in the OFC. Thus, we investigated the cellular actions of mu-opioids within distinct subregions of the OFC. Using in-vitro patch clamp electrophysiology in brain slices containing the OFC, we found that the mu-opioid agonist, DAMGO produced a concentration-dependant inhibition of GABAergic synaptic transmission onto medial OFC (mOFC), but not lateral OFC (lOFC) neurons. This effect was mediated by presynaptic mu-opioid receptor activation of local parvalbumin (PV+)-expressing interneurons. The DAMGO-induced suppression of inhibition was long-lasting and not reversed upon washout of DAMGO, or by application of the mu-opioid receptor antagonist, CTAP, suggesting an inhibitory long-term depression (iLTD) induced by an exogenous mu-opioid. We show that LTD at inhibitory synapses is dependent on downstream cAMP/PKA signaling, which differs between the mOFC and lOFC. Finally, we demonstrate that endogenous opioid release triggered via moderate physiological stimulation can induce LTD. Taken together, these results suggest that presynaptic mu-opioid stimulation of local PV+ interneurons induces a long-lasting suppression of GABAergic synaptic transmission, which depends on subregional differences in mu-opioid receptor coupling to the downstream cAMP/PKA intracellular cascade. These findings provide mechanistic insight into the opposing functional effects produced by mu-opioids within the OFC.

**Significance Statement:** Considering that both the OFC and the opioid system regulate reward, motivation, and food intake; understanding the role of opioid signaling within the OFC is fundamental for a mechanistic understanding of the sequelae for several psychiatric disorders. This study makes several novel observations. First, mu-opioids induce a long-lasting suppression of inhibitory synaptic transmission onto OFC pyramidal neurons in a regionally selective manner. Secondly, mu-opioids recruit PV+ inputs to suppress inhibitory synaptic transmission in the mOFC. Thirdly, the regional selectivity of mu-opioid action of endogenous opioids is due to the efficacy of mu-opioid receptor coupling to the downstream cAMP/PKA intracellular cascades. These experiments are the first to reveal a cellular mechanism of opioid action within the OFC.

## Introduction

The orbitofrontal cortex (OFC) plays a critical role in the evaluation of outcomes (rewards) in a changing environment. For example, the OFC is required to update actions when the value of a reward, such as food, changes with internal state (reviewed in Izquierdo, 2017). OFC neurons respond to food-predicting cues and can remain responsive after extinction of the cue-reward association, suggesting that these neurons hold long-term memories of cue-reward associations (Namboodiri et al., 2019). The OFC in rodents is located within the dorsal bank of the rostral rhinal sulcus and rostrally adjacent to the agranular insular areas, and can be subdivided into medial (mOFC), ventral (vOFC), ventrolateral (vlOFC), lateral (lOFC) and dorsolateral (dlOFC) orbitofrontal areas, identified by their proximity to the rhinal sulcus (ventral) and midline (Krettek and Price, 1977; Ray and Price, 1992; Ongür and Price, 2000). Subregions of the OFC differ based on afferent and efferent projections. Specifically, different thalamic nuclei project to the mOFC compared to the lOFC (Hunnicutt et al., 2014). Furthermore, the mOFC sends strong projections to the nucleus accumbens (NAc) core, whereas the lOFC sends significant projections to the dorsal striatum (Gremel and Costa, 2013; Murphy and Deutch, 2018). While both the mOFC and lOFC send projections to the basolateral amygdala (BLA), the mOFC to BLA projection, but not the lOFC to BLA projection is necessary for reward value retrieval (Malvaez et al., 2019), suggesting a functional segregation of these regions. Consistent with this idea, lesion studies in mice highlighted the mOFC in facilitation of goal directed response inhibition, whereas the lOFC enables acquisition of novel choices based on previous information and delays extinction of goal-directed behaviour (Gourley et al., 2010). Because there are a limited number of projections from the vOFC and mOFC to the lOFC (Hoover and Vertes, 2011), it has been proposed that the lOFC may rely less on cortico-cortical signals and instead primarily receive sensory information about cues to help guide outcome valuation (Izquierdo, 2017). These findings reveal the heterogeneity of the OFC’s afferent and efferent projections and suggest possible functional segregation between the OFC subregions.

The endogenous opioid system is also involved in the reward processing of food stimuli. Within the OFC, mu-opioid receptor availability is associated with anticipatory reward responses to palatable food (Nummenmaa et al., 2018). The mu-opioid receptor is abundantly expressed in the OFC (Mansour et al., 1995) and its signaling cascade is activated by the highly selective mu-opioid receptor peptide agonist, DAMGO. Microinjections of DAMGO into the OFC enhances both hedonic orofacial affective responses, as well as consumption of food rewards (Mucha and Iversen, 1986; Badiani et al., 1995; MacDonald et al., 2004; Mena et al., 2011; Castro and Berridge, 2017). However, there is heterogeneity in this functional mu-opioid action depending on location within the OFC. In addition to a hedonic “hotspot” found in the anterior mOFC, there is an oppositely valenced “coldspot” in the posterior lOFC, such that mu-opioid application suppresses hedonic orofacial reactions to sucrose (Castro and Berridge, 2017). These anatomically distinct functional effects suggest there are sub-regional variations in mu-opioid receptor action across the OFC. However, the cellular mechanisms underlying these differential effects by mu-opioids are unknown.

The excitability of pyramidal neurons is importantly modulated by local GABAergic interneurons, where mu-opioids act via an indirect process of disinhibition to excite these principal neurons (Zieglgänsberger et al., 1979; Madison and Nicoll, 1988; McQuiston and Saggau, 2003). While DAMGO suppresses inhibitory synaptic transmission in the vlOFC (Qu et al., 2015), its precise cellular mechanism of action across other OFC subregions is unknown. Given the distinct anatomical and functional differences between subregions of the OFC, we hypothesized that mu-opioids affect inhibitory synaptic transmission within the OFC in a regionally selective manner. To test this, we used *in-vitro* patch clamp electrophysiology to investigate the cellular actions of mu-opioids on inhibitory GABAergic transmission within the medial and lateral OFC.

## Materials and Methods

### Animals

All protocols were in accordance with the ethical guidelines established by the Canadian Council for Animal Care and were approved by the University of Calgary Animal Care Committee. Male C57BL6 mice were obtained from (Charles River Laboratories and housed in groups of 3-5). Male and female Parvalbumin (PV)-cre mouse line (JAX #: 017320 | B6 PV^cre^) were obtained from the Zamponi lab (University of Calgary Clara Christie Center for Mouse Genomics). Mice were maintained on a 12-hour light: dark schedule (lights on at 8 am MST) and were given food and water ad libitum. All experiments were performed during the animals’ light cycle.

### Electrophysiology

All electrophysiological recordings were performed in slice preparations from either male C57BL6 mice or male/female PV-cre mice (postnatal day 60-90). Briefly, mice were anaesthetized with isoflurane and were intracardially perfused with ice-cold, N-methyl D-glucamine (NMDG) solution of the following composition (in mM): 93 NMDG, 2.5 KCl, 1.2 NaH_2_PO_4_.H_2_O,30 NaHCO_3_, 20 HEPES, 25 D-glucose, 5 sodium ascorbate, 3 sodium pyruvate, 2 thiourea, 10 MgSO_4_.7H_2_O, 005 CaCl_2_.2H_2_O. Mice were then decapitated, their brain extracted, and coronal sections (250 μm) containing OFC were cut in ice-cold NMDG solution using a vibratome (Leica, Nussloch, Germany). Slices were then allowed to recover in warm NMDG solution (32°C) for 10 min, before being transferred to a long-term holding chamber containing artificial cerebrospinal fluid (ACSF) of the following composition: 126 NaCl, 1.6 KCl, 1.1 NaH_2_PO_4_, 1.4 MgCl_2_, 2.4 CaCl_2_, 26 NaHCO_3_, 11 glucose (32 °C - 34 °C). All solutions were saturated with 95% O2 / 5% CO_2_. Prior to recording, slices were transferred to a chamber on an upright microscope (Olympus BX51WI) containing ACSF, and cells were visualized using Dodt gradient contrast microscopy. Pyramidal neurons within the superficial layers of the prefrontal cortex are involved in processing long-range inputs from other regions, including the midline thalamus, basolateral amygdala and ventral hippocampus (Little and Carter, 2012). Layer II/III pyramidal neurons are particularly well positioned to integrate these synaptic inputs. Therefore, we recorded layer II/III pyramidal neurons from the mOFC or lOFC. For recordings from the mOFC, only neurons located ∼ 100-300 μm lateral to the midline and 100-500 μm dorsal to the rhinal sulcus were considered. For recordings from the lOFC, only neurons located ∼ 100-300 μm dorsal to the rhinal sulcus and ∼ 1100-1500 μm lateral from the midline were considered. Layer II/III pyramidal neurons were identified by morphological characteristics of large soma size and triangular shaped appearance, as well as electrophysiological properties of high capacitance and low input resistance. Whole-cell voltage-clamp recordings of neurons were made using a MultiClamp 700B amplifier (Axon Instruments, Union City, CA). Recording electrodes (3 - 5 MΩ) were filled with a high chloride, cesium-based internal solution of the following composition (in mM): 80 CsCH_3_SO_3_, 60 CsCl, 10 Hepes, 0.2 EGTA, 1 MgCl_2_, 2 MgATP, 0.3 NaGTP, 5 QX-314-Cl (pH 7.2–7.3, ∼ 285-290 mOsm). Series resistance (10-25 MΩ) and input resistance were monitored online with a −10 mV depolarizing step (400 ms) given before every afferent stimulus.

Neurons were voltage clamped at −70 mV and inhibitory GABA_A_-mediated IPSCs were pharmacologically isolated with the AMPA/kainate receptor antagonist, DNQX (10 μM) and the glycine receptor antagonist, strychnine (1 μM). In the majority of experiments, electrically-evoked IPSCs from intralaminar afferents were elicited at 0.1 Hz using a bipolar tungsten stimulating electrode placed either 100-300 μm dorsal of the recorded mOFC neuron, or 100-300 μm lateral of the recorded lOFC neuron. In some experiments, optically-evoked inhibitory postsynaptic currents (oIPSCs) were elicited using a 470 nM LED light flash (4-5 mW, Thorlabs) via the microscope objective. IPSCs were filtered at 2 kHz, digitized at 10 kHz and collected using pCLAMP 10 software. In other experiments, miniature IPSCs (mIPSCs) were recorded in the presence of the Na+ channel blocker, tetrodotoxin (500 nM) to isolate non-action potential dependent spontaneous release. mIPSCs were selected for amplitude (> 15 pA), decay time (< 10 ms) and rise time (< 4 ms) using the MiniAnalysis program (Synaptosoft).

### Immunohistochemistry and Confocal Microscopy

Brains were perfused and stored overnight in 4% paraformaldehyde, then switched to 20% sucrose and coronal frozen sections were cut at 30 μm using a cryostat. Sections were then blocked in serum before incubation with mouse monoclonal β-endorphin primary antibody (Abeam ab54205,1:200) for 24 h at 4°C and Alexa-fluor 488 chicken anti-mouse secondary antibody (Invitrogen, 1:400) for 24 hours at 4°C, and slices were mounted with Fluoromount. In some cases, OFC β-endorphin sections were co-labeled with the primary antibody for goat anti parvalbumin (PV+) (1:1000, SIGMA, SAB2500752), a marker for a cortical interneuron subtype, in blocking solution for 24 h at 4°C. After washing, secondary antibody was applied (Alexa fluor-594 donkey anti goat IgG; 1:400) in blocking solution for 3 hrs. Slices were then washed and mounted with DAPI. For met-enkephalin staining, OFC sections were reacted with the primary antibody for rabbit met-enkephalin (1:400, Abeam ab22620) for 24h at 4°C and Alexa-fluor 488 goat anti rabbit secondary antibody (Invitrogen, 1:400) for 24 hours at 4°C. 6 slices taken from 3-4 mice per condition were imaged. All images were obtained on a Nikon Eclipse C1si confocal microscope with a motorized stage (Nikon Canada Inc. Ontario, Canada). The objectives used were 20X Plan Apo DIC (NA 0.75).

### Stereotaxic Surgery

PV-cre mice were anesthetised with isoflurane and placed in a stereotaxic frame (David Kopf Instruments; Tujunga, CA, USA). AAV2-EF1a-DIO-hChR2(H134R)-EYFP (Neurophotonics; lot 227, 4.3 x 10^12^ GC / mL) was injected into the mOFC (Anterior/posterior: + 2.68, mediolateral: ± 0.55, dorsoventral: 200 nl @ −1.945, 200 nl @ −1.445). A total of 400 nl of virus was injected in each hemisphere at 23 nl/s using the Nanoject II (Drummond Scientific Company) allowing 3 minutes for the virus to disperse throughout the tissue following each injection. Mice were treated post-surgically with ketoprofen (5 mg/kg, subcutaneous) and were left for at least 3 weeks post-surgery to allow for expression of ChR2 and EYFP in PV+ neurons of the mOFC.

### Statistics

Data are expressed as mean ± SEM. Data were analyzed with a paired t-test (for comparing before and after conditions) or repeated measures one-way ANOVA for measuring drug effects and wash in the same neuron, unless otherwise indicated. Unless otherwise indicated, data met the assumptions of equal variance. In all electrophysiology experiments, sample size is expressed as n/N where “n” refers to the number of cells recorded from “N” animals. Graph Pad Prism 7 software (GraphPad Software, Inc., La Jolla, CA) was used to perform statistical analysis. The levels of significance are indicated as follows: *, **, *** denote P < 0.05, 0.01, 0.001.

## Results

### Mu-opioid receptor activation has no effect on GABAergic synaptic transmission onto lOFC pyramidal neurons

Given that mu-opioids elicit regionally distinct orofacial affective responses in the OFC (Castro and Berridge, 2017), and that they act predominantly via disinhibition to excite output neurons, we hypothesized that mu-opioids differentially modulate inhibitory synaptic transmission onto pyramidal neurons within the mOFC and lOFC. We first examined the effect of the selective mu-opioid receptor agonist, DAMGO (1 μM) on eIPSCs from layer II/III pyramidal neurons in the lOFC (Fig.1A,B). DAMGO did not significantly alter elPSC amplitude onto lOFC pyramidal neurons (baseline: 98 ± 2, DAMGO: 100 ± 4, wash: 99 ± 4; F(1.8,18.1) = 0.12, P = 0.87, n/N = 11/8; Fig. 1C-E). The paired pulse ratio, a measure that typically correlates with release probability, was not significantly different before and after DAMGO application (t_(10)_ = 0.70, P = 0.5, n/N = 11/8; Fig. 1F). Taken together, mu-opioid receptor activation does not alter inhibitory synaptic transmission onto layer II/III pyramidal neurons of the lOFC.

**Figure 1.**
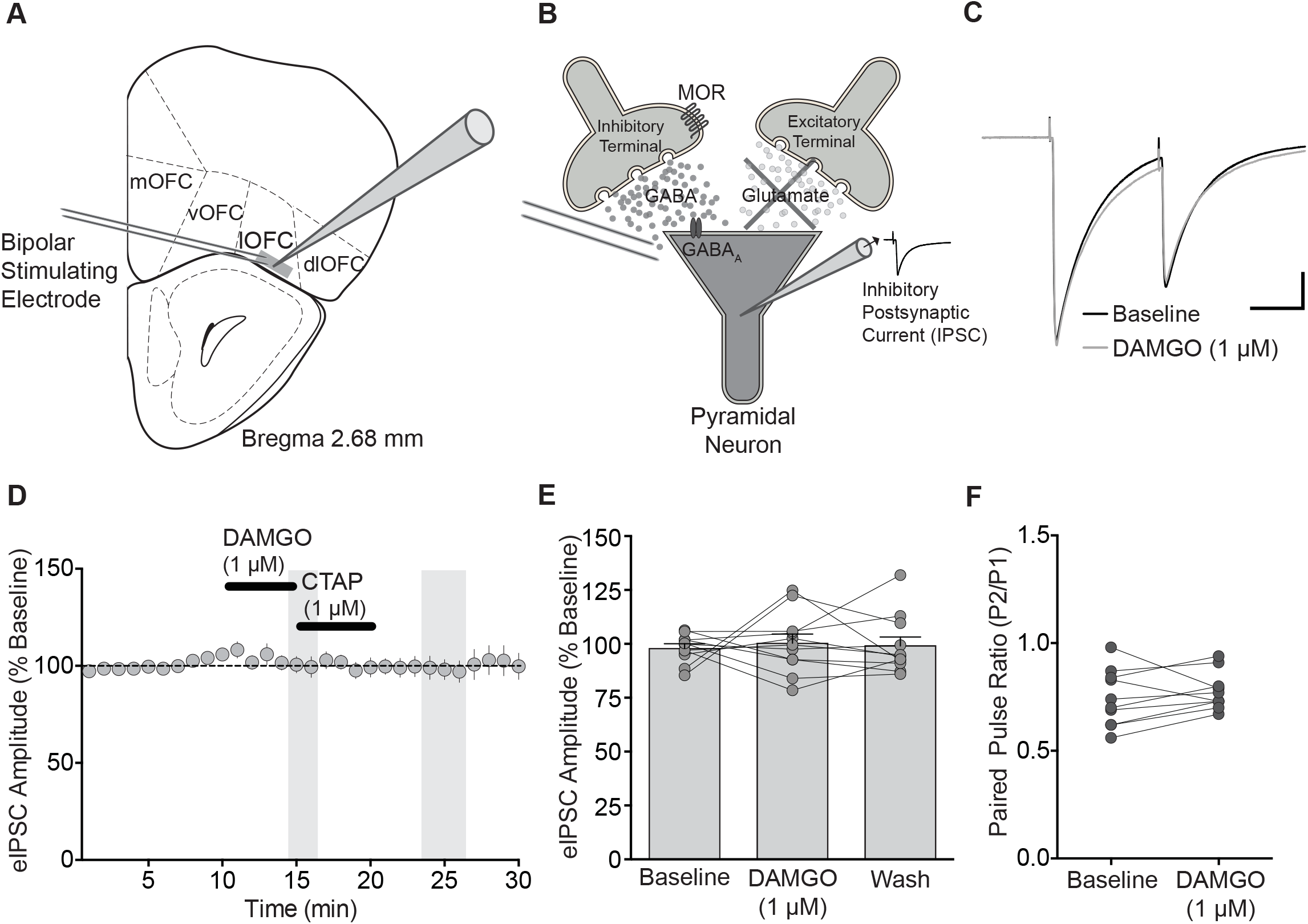
DAMGO does not alter eIPSCs in layer II/III of the lOFC. **A.** Schematic indicating the lateral subregion of the orbitofrontal cortex where inhibitory synaptic transmission onto layer II/III pyramidal neurons was recorded (schematic modified from Paxinos and Franklin, 2001, Bregma −2.68 mm). **B.** Schematic indicating locally-evoked, pharmacologically isolated IPSCs onto recorded pyramidal neuron. C. Averaged example traces of elPSCs in lOFC neurons before (Baseline) and after application of DAMGO (1 μM, 5 min). Paired pulses are 50 ms apart. Scale bar: 250 pA, 25 ms. D. Time course of eIPSC before and during DAMGO application and then washout with the selective mu-opioid receptor antagonist, CTAP (1 μM). Symbols represent averaged responses ± SEM per minute, n/N = 11/9. Shaded bars represent time periods during DAMGO and after washout which were analyzed in of E. DAMGO had no effect on elPSC amplitude in lOFC pyramidal neurons. E. Scatter plot quantifying individual responses before and after DAMGO application. F. Scatter plot of individual paired-pulse ratio before and after DAMGO application. Bars represent mean.

### Mu-opioid receptor activation inhibits GABAergic synaptic transmission onto mOFC pyramidal neurons

We next examined the effect of DAMGO on GABAergic synaptic transmission within the mOFC (Fig. 2A,B). In contrast to the lOFC, DAMGO (1 μM) produced an irreversible inhibition of eIPSC amplitude in mOFC pyramidal neurons (baseline: 104 ± 3%, DAMGO: 59 ± 8%, wash: 50 ± 4%; F(1.4, 8.4) = 23.24, P = 0.0007, Tukey’s multiple comparison test indicates significant differences between baseline and DAMGO (p <0.05) and baseline and wash (p<0.0001), n/N = 7/5; Fig. 2C-E). Notably, the selective mu-opioid receptor antagonist, CTAP (1 μM) did not reverse DAMGO-mediated inhibition of eIPSC amplitude, suggesting that mechanisms downstream of mu-opioid receptor activation may be required to maintain the synaptic depression. Paired pulse ratio was not significantly different after DAMGO application (t_(6)_ = 1.86, P = 0.1, n/N = 7/5; Fig. 2F). These results indicate that mu-opioid receptor activation decreases inhibitory synaptic transmission onto layer II/III pyramidal neurons of the mOFC.

**Figure 2.**
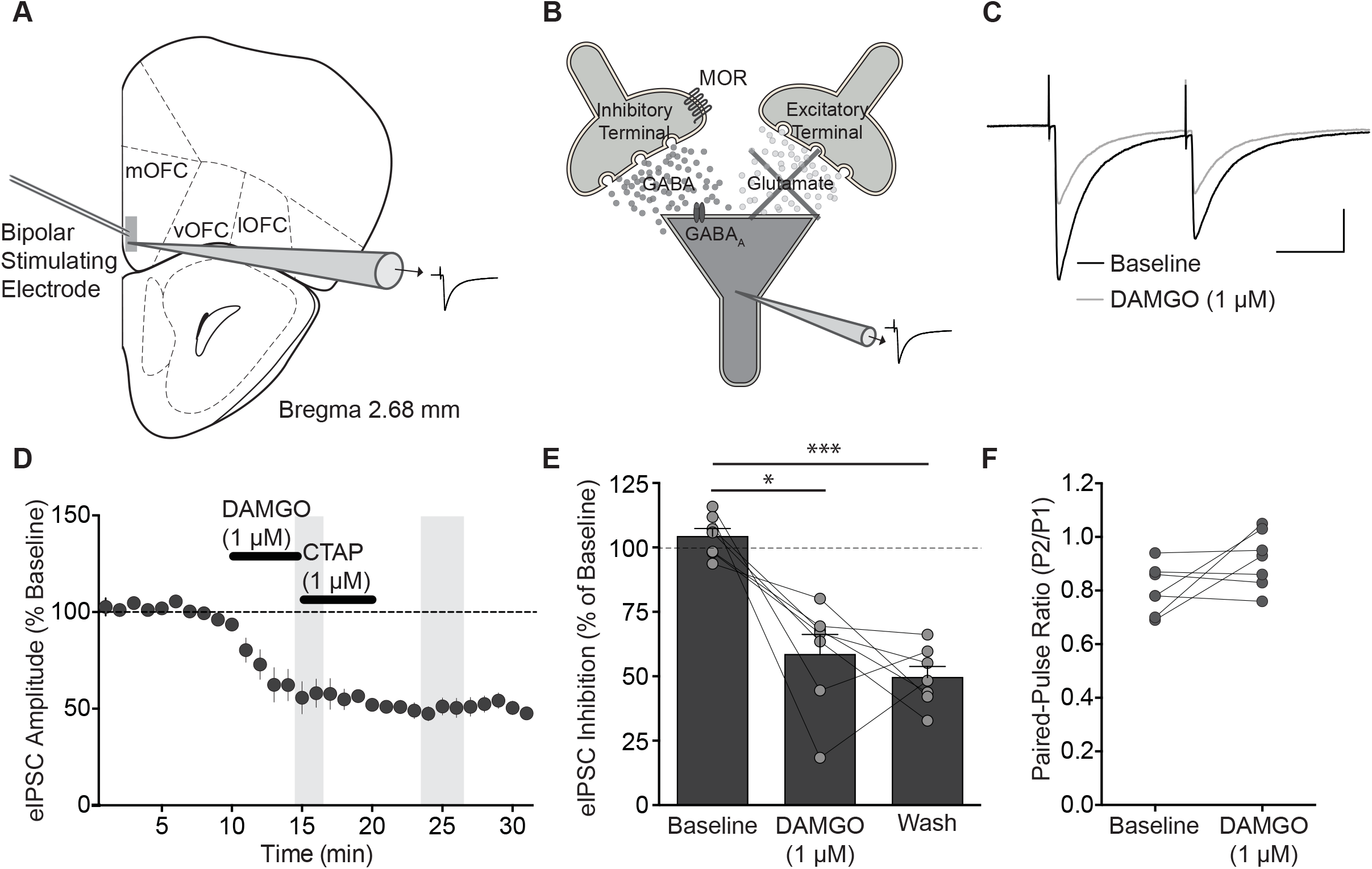
DAMGO induces a long-lasting suppression of eIPSCs onto layer II/III pyramidal neurons of the mOFC. **A.** Schematic indicating the medial subregion of the orbitofrontal cortex where inhibitory synaptic transmission onto layer II/III pyramidal neurons was recorded (Schematic modified from Paxinos and Franklin 2001, Bregma −2.68 mm). B. Schematic illustrating the recording of evoked IPSCs onto a mOFC pyramidal neuron, which were elicited via intralaminar electrical stimulation and pharmacologically isolated with the AMPA/kainate receptor antagonist, DNQX (10 μM). C. Averaged example traces of eIPSCs in mOFC neurons before and after DAMGO (1 μM) application. Paired pulses are 50 ms apart. Scale bar, 250 pA, 25 ms. D. Time course of eIPSCs before and after a 5 min DAMGO application with the selective mu-opioid antagonist, CTAP (1 μM). Symbols represent averaged responses ± s.e.m. n/N = 7/5. Shaded bars represent time periods during DAMGO and after washout which were analyzed in of E. DAMGO produced irreversible suppression of eIPSC amplitude in mOFC pyramidal neurons. E. Individual responses before and after DAMGO application (*, P<0.05, **P<0.01). **F.** Paired pulse ratio measured from individual cells before and after DAMGO application.

To confirm that the DAMGO-induced suppression of eIPSCs in the mOFC was mediated by mu-opioid receptors, we bath applied CTAP (1 μM) for 10 min prior to DAMGO (1 μM, 5 min) application (Fig. 3A,B). Pre-application of CTAP did not affect baseline eIPSC amplitude, but blocked the DAMGO-mediated inhibition of eIPSCs, indicating that this effect requires activation of mu-opioid receptors (baseline + CTAP: 99 ± 2%, DAMGO + CTAP: 92 ± 5%, CTAP Wash: 98 ± 5%; F(1.19, 4.57) = 0.79, P = 0.439, n/N = 5/4; Fig. 3A,B). We next tested the concentration-dependence of this effect by DAMGO (Fig. 3C). DAMGO (100 nM) irreversibly inhibited eIPSC amplitude to a maximum of 40±7% (baseline: 101 ± 3%, DAMGO: 61 ± 5%, wash: 80 ± 5, F(1.91, 11.47) = 16.01, P = 0.0005, n/N = 7/s)o Because the magnitude of this inhibition was indifferent from 1 μM, 100 nM DAMGO is a saturating concentration for inhibition of eIPSC in the mOFC. DAMGO (10 nM) significantly inhibited eIPSCs by 15 ± 3 %, and washout of DAMGO or subsequent application of CTAP did not reverse the effect of this submaximal DAMGO concentration (baseline: 97 ± 1, DAMGO: 82 ± 2, wash: 83 ± 4; F(1.50, 11.96) = 7.88, P = 0.0097). ATukey’s multiple comparison test indicates significant differences between baseline vs. DAMGO (p=0.0021) and baseline vs CTAP (P = 0.029); n/N = 9/7; Fig. 3D, E). Paired-pulse ratio was not significantly different after DAMGO (10 nM) (t_(8)_ = 1.844, P = 0.10, n/N = 9/7; Fig. 3F). Finally, 1 nM DAMGO did not significantly suppress eIPSCs onto mOFC pyramidal neurons (baseline: 101 ± 2, DAMGO: 94 ± 5; CTAP: 89 ± 5; F(1.63, 6.51) = 1.92, P = 0.21, n/N = 5/5). A logarithmic dose response curve calculated a half-maximal inhibition concentration (IC_50_) of 18 nM DAMGO on eIPSCs of mOFC pyramidal neurons (Fig. 3C).

**Figure 3.**
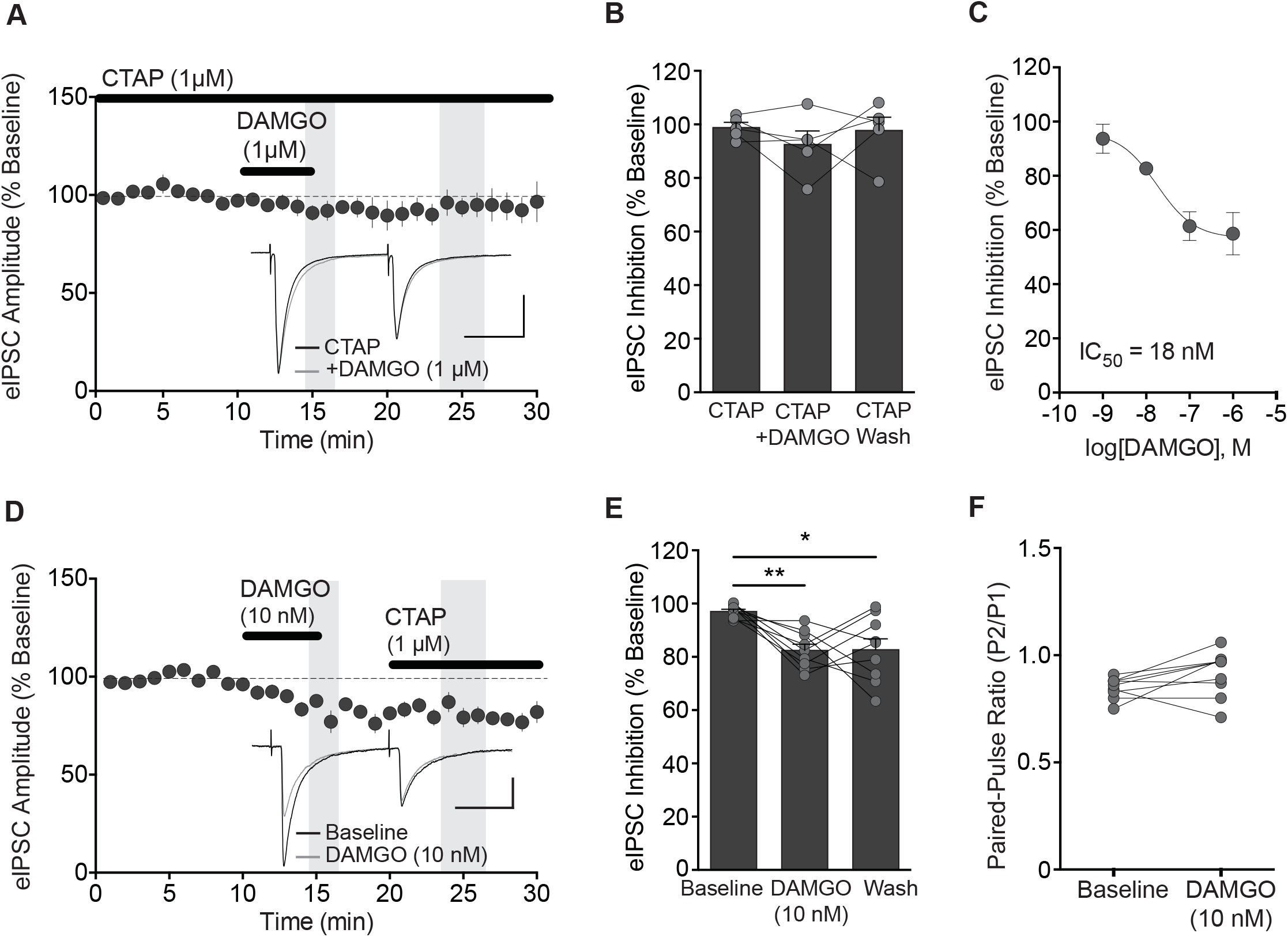
DAMGO concentration dependently inhibits eIPSC via mu-opioid receptors. **A.** Time course of eIPSC amplitude showing the effect of DAMGO (1 μM) on mOFC neurons in slices pre-treated with CTAP (1 μM). Shaded bars represent time periods during DAMGO and after washout which were analyzed in of B. In the presence of CTAP, a 5 min application of DAMGO does not alter eIPSCs onto layer II/III mOFC neurons (n/N= 6/4). Scale bars: 250 pA, 25 ms. Inset: Averaged sample traces of eIPSCs before and after DAMGO application in the presence of CTAP. B. Bar graph quantifying the eIPSC amplitude before, during and after DAMGO application in the presence of CTAP. **C.** Concentration-response curve fit with a sigmoidal curve showing a maximal inhibition at 100 nM and an IC_50_ of 18 nM. N = 6-9 cells per concentration. D. Time course of eIPSC amplitude showing the effect of DAMGO (10 nM) in mOFC neurons. Shaded bars represent time periods during DAMGO and after washout with CTAP which were analyzed in E. A lower concentration of DAMGO (10 nM) induces a suppression of eIPSC amplitude which is not reversed by CTAP (1 μM) application. n/N = 9/7, Inset: Averaged traces of eIPSCs before and during DAMGO (10 nM) application, Scale bar: 100 pA, 25 ms. E. Bar graph quantifying the eIPSC amplitude before and during application of DAMGO, and then following washout/CTAP application. * and ** denote P < 0.05 and 0.01. F. Bar graph quantifying paired-pulse ratio before and during DAMGO application.

To determine whether the effect of DAMGO on inhibitory transmission was mediated by a presynaptic or postsynaptic mechanism, we recorded GABA_A_-mediated mIPSCs onto mOFC pyramidal neurons. While DAMGO (1 μM, 5 min) did not alter the amplitude of mIPSCs (Fig. 4A,B; baseline: 35 ± 1.7 pA; DAMGO: 34 ± 1.4 pA; t(7) = 0.81, P = 0.4, n/N = 8/3), DAMGO significantly decreased the frequency of mIPSCs (baseline: 8.5 ± 0.5 Hz; DAMGO: 6.9 ± 0.4 Hz; t(7) = 3.67, P = 0.0079, n/N = 8/3; Fig. 4A,C). The maximum difference between the cumulative distributions of mIPSC amplitude in baseline or DAMGO was not significantly different (Kolmorgorov-Smirnov (K-S) test: D_(111)_ = 0.0982, *P* = 0.630; Fig. 4B). However, there was a significant difference in cumulative distribution for mIPSC frequency (D_(111)_= 0.5446, *P* < 0.0001; Fig. 4C). In contrast, DAMGO (1 μM) did not significantly alter mIPSC amplitude (baseline: 34 ± 1.7 pA; DAMGO: 35 ± 1.2 pA; t(14) = 1.21, P = 0.2, n/N = 15/4) or frequency (baseline: 9 ± 0.8 Hz; DAMGO: 9 ± 0.8 Hz; t(14) = 0.82, P = 0.4, n/N = 15/4) in the presence of CTAP (1 μM; Fig. 4D,E, F). Furthermore, there was no significant difference in cumulative distributions during CTAP between baseline or DAMGO amplitude (D_(101)_ *=* 0.0536, P = 0.996; Fig. 4E) or frequency (D_(101)_ *= 0*.1485, P = 0.198; Fig.4F). Taken together, mu-opioid receptor activation likely suppresses GABA release onto mOFC pyramidal neurons.

**Figure 4.**
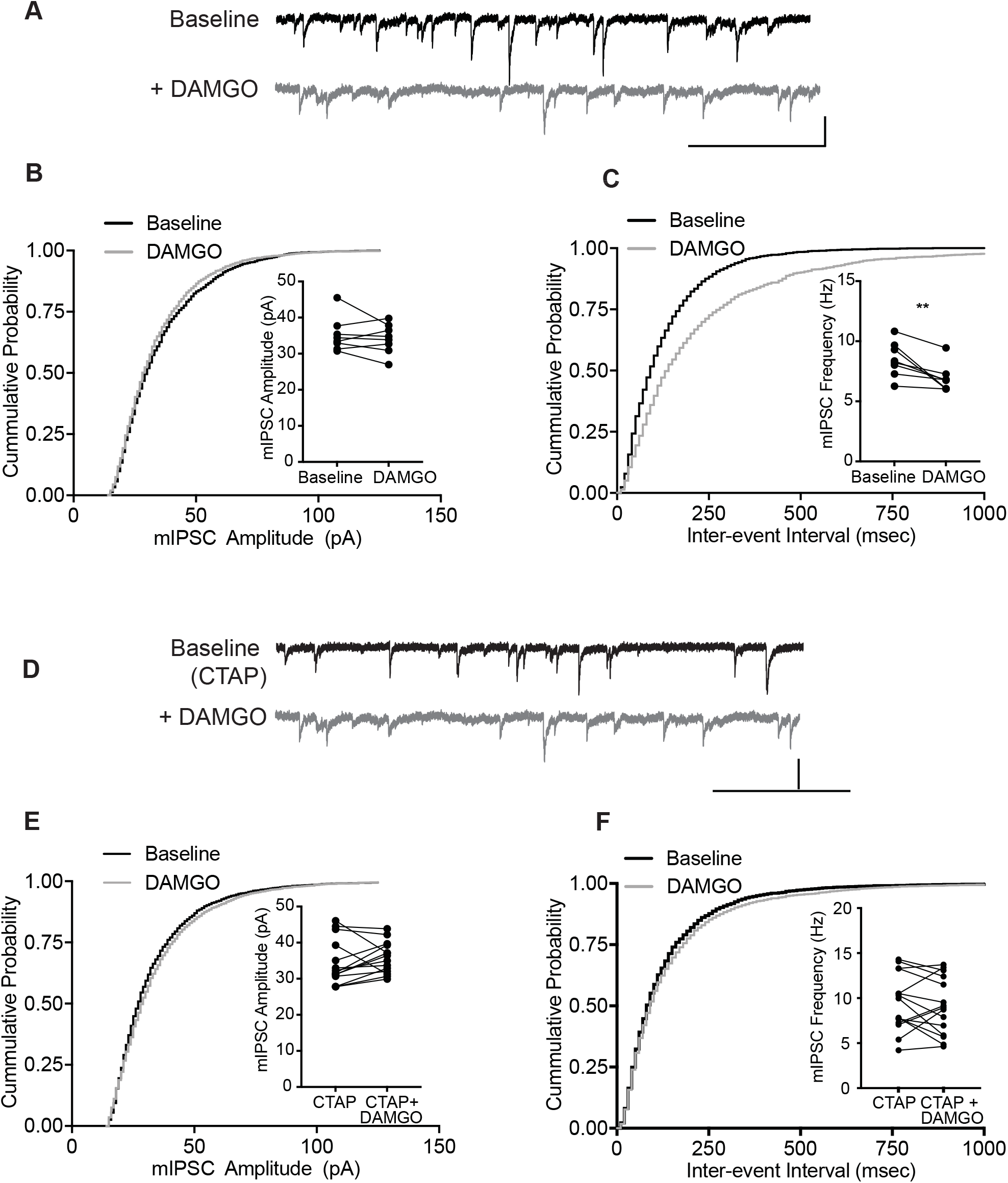
DAMGO suppresses GABA release. **A.** Representative example traces of mIPSCs from before (Baseline) and during DAMGO (ιμM). Scale bars, 25 pA, 250 ms. B. Cumulative distribution probability curve for mIPSC amplitude before and after DAMGO (1μM) application. Inset, averaged mIPSC event amplitude of individual mOFC neurons before and during DAMGO, n/N = 8/3. C. Cumulative distribution probability curve for mIPSC inter-event intervals before and after DAMGO (1 μM) application. Inset, averaged mIPSC frequency of individual mOFC neurons before and after DAMGO (1μM) application (**p<0.01). n/N = 8/3. D. Representative example traces of mIPSCs from before (Baseline) and during DAMGO (1μM) in the presence of CTAP (1 μM). Scale bars: 25 pA, 250 ms. E. Cumulative distribution probability curve for mIPSC amplitude before and during DAMGO (1 μM) in the presence of CTAP (1 μM). Inset, averaged mIPSC event amplitude of individual mOFC neurons before and after DAMGO in the presence of CTAP. n/N = 15/4. F. Cumulative distribution probability curve for mIPSC inter-event intervals before and during DAMGO (1μM) in the presence of CTAP (1 μM). Inset, averaged mIPSC frequency of individual mOFC neurons before and during DAMGO in the presence of CTAP. n/N = 15/4.

### Mu-opioids target PV+ terminals to suppress GABA release onto mOFC pyramidal neurons

A variety of fast-spiking interneurons, including parvalbumin expressing (PV+) interneurons, release GABA onto OFC pyramidal neurons (Quirk et al., 2009). PV+ interneurons play a significant role in coordinating gamma oscillations in the cortex to influence cortical network activity (Sohal et al., 2009). Mu-opioid receptors are expressed on PV+ interneurons in the dentate gyrus (Drake and Milner, 2006) as well as somatosensory and frontoparietal cortices (Taki et al., 2000). To determine whether DAMGO influences inhibitory synaptic transmission from PV+ interneurons onto mOFC pyramidal neurons, a cre-recombinase dependent adeno-associated virus (AAV) virus expressing ChR2 was administered into the mOFC of PV-cre mice (Fig. 5A). Following intra-mOFC AAV-ChR2 injection, YFP-staining was found in both cell bodies and puncta throughout the mOFC (Fig. 5B). Photostimulation (5 mW, 0.1 ms, 470 nm LED) of PV+ terminals readily evoked IPSCs onto mOFC pyramidal neurons (Fig. 5C). DAMGO (1 μM, 5 min) significantly decreased optically-evoked IPSC (oIPSC) amplitude to 57 ± 6% of baseline (F(1.36,6.82) = 17.83, P = 0.0029, n/N = 10/6; Fig. 5C-E). Consistent with mu-opioid receptor effects on electrically evoked IPSCs, this suppression of inhibition was long-lasting and was not reversed by washout of DAMGO or subsequent addition of CTAP (Fig. 5D). A Tukey’s multiple comparison test indicated a significant difference between baseline and DAMGO (P = 0.014) and baseline and wash (P = 0.0163). By contrast, the DAMGO-induced suppression of oIPSC amplitude was blocked in slices pre-treated with CTAP (1 μM, 100 ± 3%, n/N = 6/4, F(1.55, 7.748) = 0.37, P = 0.65; Fig. 5D,E). We additionally examined the effect of DAMGO on ChR2-expressing, PV-inputs onto lOFC neurons. Similar to the effects on eIPSCs, DAMGO had no effect on olPSC amplitude in lOFC neurons (F(1.3, 7.7) = 1.28, P > 0.05; Fig. 5F-H). Together, these data support the hypothesis that activation of mu-opioid receptors on PV+ interneurons suppresses GABAergic transmission onto mOFC, but not lOFC pyramidal neurons.

**Figure 5.**
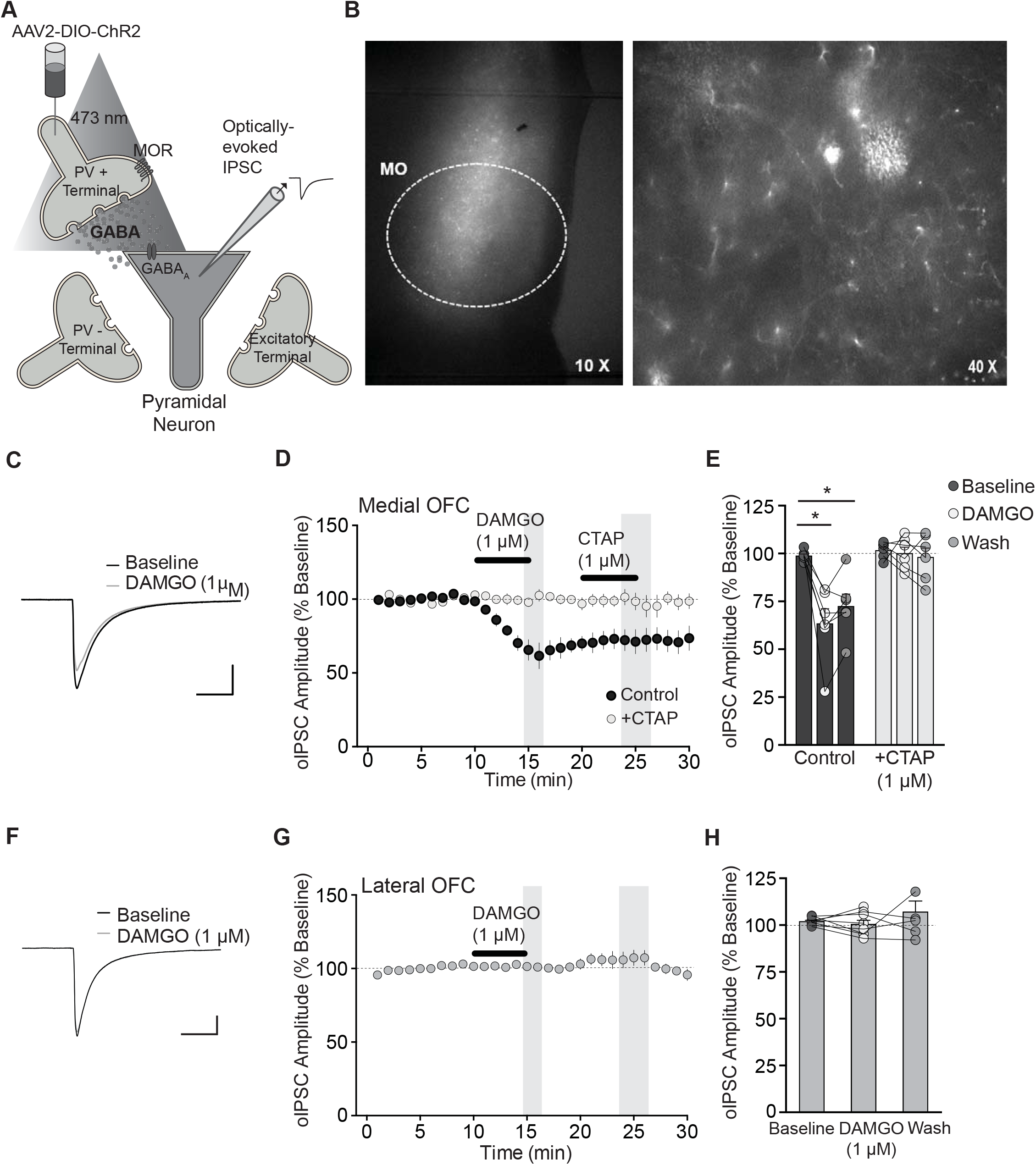
DAMGO induces a long-lasting suppression of oIPSCs from PV+ inputs to pyramidal neurons. **A.** Schematic illustrating optical stimulation of ChR2 expressed in local PV neurons and recording their inputs onto pyramidal neurons within the mOFC. **B.** Low (10X) and high (40X) magnification images of ChR2 expresssion in PV+ neurons within the mOFC. **C**. Representative traces of oIPSCs onto mOFC pyramidal neurons elicited via optical stimulation of PV inputs before and during DAMGO (1 μM) application. Scale bars, 250 pA, 25 ms. **D.** Time-course of oIPSC amplitude before, during and after DAMGO (1 μM) application (filled symbols, n/N = 6/4) or in the presence of CTAP (1 μM) (open symbols, n/N = 6/4). Symbols represent mean ± SEM. **E**. Bar graph with individual responses quantifying DAMGO inhibition of oIPSC amplitude in mOFC neurons from control slices or those pre-treated with CTAP. **F.** Representative traces of olPSCs onto lOFC pyramidal neurons elicited via optical stimulation of PV inputs before and after DAMGO (1 μM) application. Scale bars, 250 pA, 25 ms. **G.** Time-course of oIPSC amplitude before, during and after DAMGO (1μM) application. DAMGO had no effect on olPSC amplitude in lOFC neurons. n/N = 8/5 Symbols represent averaged responses ± SEM per minute. **H.** Bar graph quantifying oIPSC amplitude before and during application of DAMGO (1 μM).

### Downstream cAMP/PKA intracellular signalling mediates mu-opioid long-term depression of inhibitory transmission

We next tested a potential mechanism for the long-lasting effects of DAMGO-induced LTD in the mOFC. Activation of mu-opioid receptors typically results in a G_i/o_-protein mediated inhibition of adenylyl cyclase, which in turn catalyzes the conversion of ATP into cyclic adenosine monophosphate (cAMP), leading to activation of PKA. Both cAMP and PKA are required for many forms of presysnaptic long-term depression (LTD) (Mato et al., 2008; Yang and Calakos, 2013). To explore the role of the cAMP/PKA cascade in mediating mu-opioid-mediated LTD in the mOFC, we examined the effect of the cAMP/PKA activator, forskolin (10 μM) on eIPSCs. Application of forskolin to mOFC neurons induced a facilitation of eIPSC amplitude (130 ± 6% of baseline, n/N = 9/8, one-sampled t-test to 100% baseline: t_(9)_ = 4.98, P = 0.0008; Fig. 6A-C). Subsequent addition of DAMGO significantly reduced eIPSC amplitude (85 ± 4% of forskolin, n/N = 9/8, one sampled t-test: t_(9)_ = 3.53, P = 0.0064; Fig. 6A,B,D). However, this suppression of inhibition was reversed upon washout (105 ± 5% of forskolin, N = 9/8, t_(9)_ = 0.10, P = 0.34; Fig. 6D-F). In contrast, application of forskolin to lOFC neurons had no effect on eIPSC amplitude (105 ± 7% of baseline, n/N = 5/4, t_(6_) = 0.69, P = 0.52; Fig. 6A-C) and this response was unchanged by subsequent addition of DAMGO or upon washout (DAMGO: 103 ± 9% of forskolin, t_(5)_ = 0.09, P = 0.93, washout: 109 ± 9% of forskolin, t_(5)_ = 1.60, P = 0.17, n/N = 5/4; Fig. 6A,B,D). The effect of forskolin on eIPSC amplitude was significantly greater in mOFC neurons than those of the lOFC (unpaired t-test: t(15) = 2.815, P = 0.0131; Fig. 6C). Furthermore, there was a significant interaction of OFC subregion and drug treatment on eIPSC amplitude (2 way RM ANOVA: F(1,14) = 6.491, P = 0.023) and a main effect of drug treatment (F(1,14) = 31.17, P < 0.001). A Sidak’s multiple comparisons test indicated a significant difference between DAMGO and wash in the mOFC (P<0.0001), but not lOFC (P>0.05; Fig. 6D). Together, these results indicate that there is differential coupling of mu-opioid receptors to cAMP/PKA between mOFC and lOFC pyramidal neurons, and that cAMP/PKA signalling is required for the maintenance of mu-opioid-mediated LTD in mOFC neurons.

**Figure 6.**
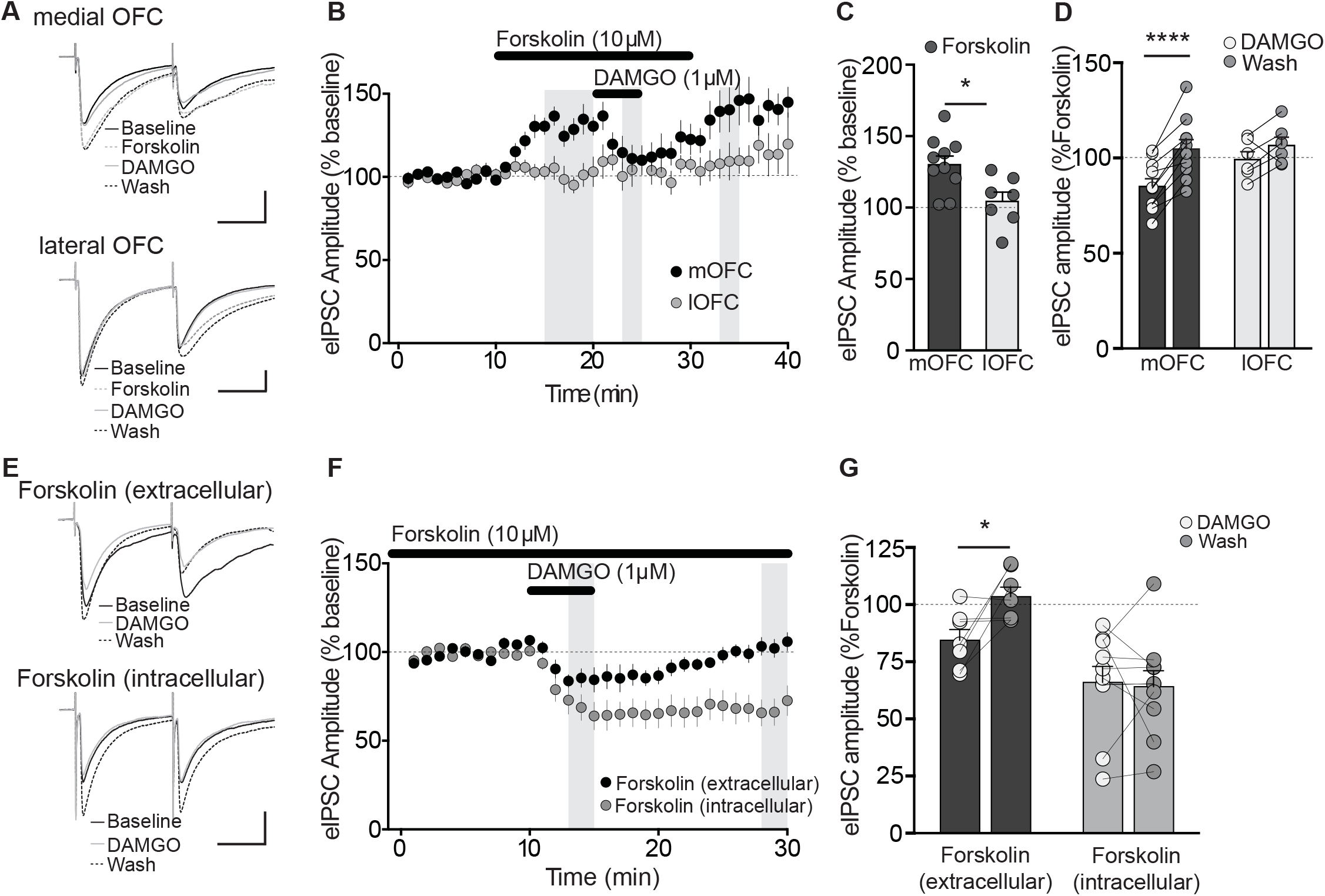
Mu-opioid receptors require downstream cAMP/PKA signalling to induce long-term depression of inhibitory transmission. **A.** Example traces of eIPSCs before (baseline) and during application of forskolin (10 μM), plus further addition of DAMGO (10 μM), then after washout of these drugs in mOFC or lOFC neurons. Scale bars: 250 pA, 100 ms. B. Time course of eIPSC amplitude before and during application of forskolin (10 μM), DAMGO, and then washout in mOFC or lOFC neurons. Shaded bars represent time periods analyzed in bar graphs of C and D. Forskolin induces a facilitation of eIPSC amplitude in mOFC, but not lOFC neurons. Subsequent addition of DAMGO reversibly depressed eIPSC amplitude in mOFC neurons, but had no effect in lOFC neurons. C. Bar graph quantifying the percentage change in eIPSC amplitude produced by forskolin relative to baseline. There was a significant difference in forskolin response between neurons of mOFC and lOFC (*P<0.05). D. Bar graph and individual responses quantifying the response to DAMGO and washout relative to forskolin. A Sidak’s posthoc test indicates a significant difference between DAMGO and washout for mOFC (****P<0.0001), but not lOFC neurons (P>0.05). E. Example traces of eIPSCs before, during and after DAMGO application in mOFC neurons, in which forskolin was present in the extracellular ACSF or intracellular pipette solution. Scale bars: 200 pA, 25 ms. F. Time course of eIPSC amplitude before, during and after DAMGO application in mOFC neurons, in which forskolin was either present in the extracellular ACSF (filled symbols) or intracellular pipette solution. Shaded bars represent time periods analyzed in bar graphs of G. DAMGO induced a reversible short-term suppression of eIPSC amplitude with intracellular forskolin application, but an irreversible long-term suppression with extracellular forskolin application. G. Bar graph quantifying the change in eIPSC amplitude during and after DAMGO application in mOFC neurons pre-treated with either extracellular or intracellular forskolin. A Sidak’s multiple comparison test indicates a significant difference between DAMGO and wash during intracellular (*P<0.05), but not extracellular forskolin application (P>0.05).

To further confirm the locus of this cAMP/PKA-mediated effect in the mOFC, we pre-applied forskolin (> 20 min) either intracellularly in the recording pipette to confine its action postsynaptically in the recorded neuron, or extracellularly in the ACSF bath to act presynaptically and postsynaptically. Pyramidal neurons from the mOFC pre-treated with forskolin in the extracellular ACSF exhibited a reversible suppression of eIPSC by DAMGO (DAMGO: 85 ± 5%, Wash: 104 ± 4%, n/N = 10/7; Fig. 6E-G). In contrast, mOFC neurons pre-loaded with forskolin in the intracellular pipette exhibited an irreversible suppression of eIPSCs by DAMGO (DAMGO: 66 ± 7%, Wash: 64 ± 7%, n/N = 7/7; Fig. 6E-G). There was a forskolin x DAMGO interaction (2 way RM ANOVA: F(1,15) = 4.45, P = 0.05) and a main effect of forskolin site (F(1,15) = 14.02, P = 0.002). A Sidak’s multiple comparison test indicates a significant difference between DAMGO and washout only when forskolin is applied extracellularly (P<0.05). Together, these results indicate that DAMGO-induced LTD in mOFC neurons is mediated upstream by cAMP/PKA signaling in presynaptic GABA inputs. The lack of effect by DAMGO in the lOFC is likely due to reduced functional coupling between mu-opioid receptors and their downstream intracellular cascades.

### Endogenous opioid expression is similar between the mOFC and lOFC

A possible explanation for the differential coupling of mu-opioid receptors to cAMP/PKA in the mOFC and lOFC is varying expression of endogenous opioids, which may facilitate or occlude activation of mu-opioid receptors in these subregions. To examine endogenous sources of mu-opioids, we used immunohistochemistry to determine β-endorphin and met-enkephalin expression within the lOFC and mOFC. We also assessed expression in the dorsal striatum as a positive control for met-enkephalin and β-endorphin-like immunoreactivity. Consistent with previous reports, we found strong met-enkephalin (Extended Data Fig. 7-1) and β-endorphin-like immunoreactivity in the striatum (Fig. 7C). There was no significant difference in the intensity of β-endorphin-like expression in the mOFC, lOFC, or striatum ((F(2,14) = 0.078, P = 0.93; Fig. 7A-D). We then examined whether β-endorphin was co-localized with PV-expressing neurons. We observed PV-like expression in the mOFC, lOFC and striatum, but there was no significant difference in the intensity of its expression in all three regions (F(2,17) = 0.15, P = 0.85; Fig. 7A,B,C,E). PV was only weakly co-localized with β-endorphin at similar levels in all 3 regions (11 ± 3%, 7 ± 3% and 6 ± 5% respectively, n = 9, 8, 6, respectively; F_(2,20)_ = 0.84, P = 0.44; Fig. 7F). Therefore, the above results indicate that endogenous opioids are expressed at similar levels within the mOFC and lOFC

**Figure 7.**
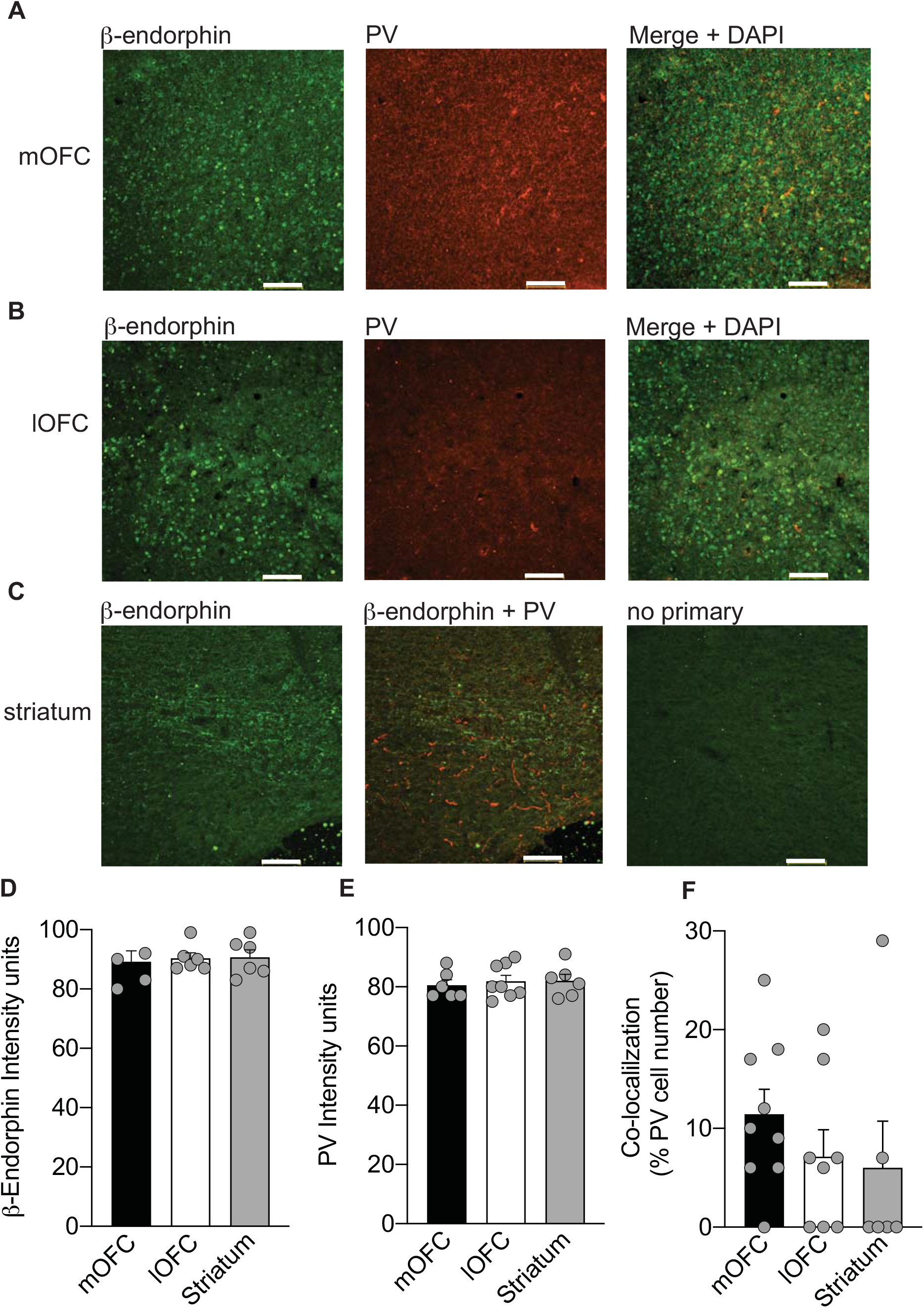
Endogenous opioids are locally expressed in mOFC and lOFC. **A.** Immunoreactivity of β-endorphin (green), PV (red) and merged image in the mOFC. Scale bar, 100 μm. **B**. Immunoreactivity of β-endorphin (green), PV (red) and merged image in the IO FC. Scale bar, 100 μm. **C**. Immunoreactivity of β-endorphin (green), β-endorphin + PV (green + red) and a no primary negative control in the striatum. Scale bar, 100 μm. **D**. Bar graph quantifying the fluorescence intensity of β-endorphin in the mOFC, lOFC and striatum. **E**. Bar graph quantifying the fluorescence intensity of PV in the mOFC, lOFC and striatum per region of interest. **F.** Bar graph quantifying the degree of co-localization between PV and β-endorphin cells (as a percentage of total PV+ neurons) in the mOFC, lOFC and striatum.

### Endogenously released opioids facilitate long-term depression of GABA synapses in the mOFC under conditions of reduced metabolic degradation

Given that endogenous opioids are present in both the mOFC and lOFC, we then tested whether they exert a functional influence on GABA synapses. If endogenous opioids are present and suppressing GABAergic transmission under basal conditions, then blockade of mu-opioid receptors by CTAP should unmask a facilitation of GABA release. Application of CTAP (1 μM) in the lOFC showed a trend towards increasing eIPSC amplitude, but this was not significantly different from baseline (112 ± 7%, n/N = 10/6, one sampled t-test to 100%; t_(9)_ = 1.822, P=0.10; Fig. 8A-C). Furthermore, CTAP application had no effect in the mOFC (95 ± 5%, n/N = 5/4, t_(4)_ = 1.058, P = 0.35; Fig. 8A-C). Overall, there was no difference in CTAP effect between mOFC and lOFC neurons (t_(13)_ = 1.69, P = 0.114). Thus, while local sources of endogenous opioids are present in the mOFC and lOFC, they do not actively induce a tone at inhibitory synapses onto mOFC or lOFC neurons under basal conditions.

**Figure 8.**
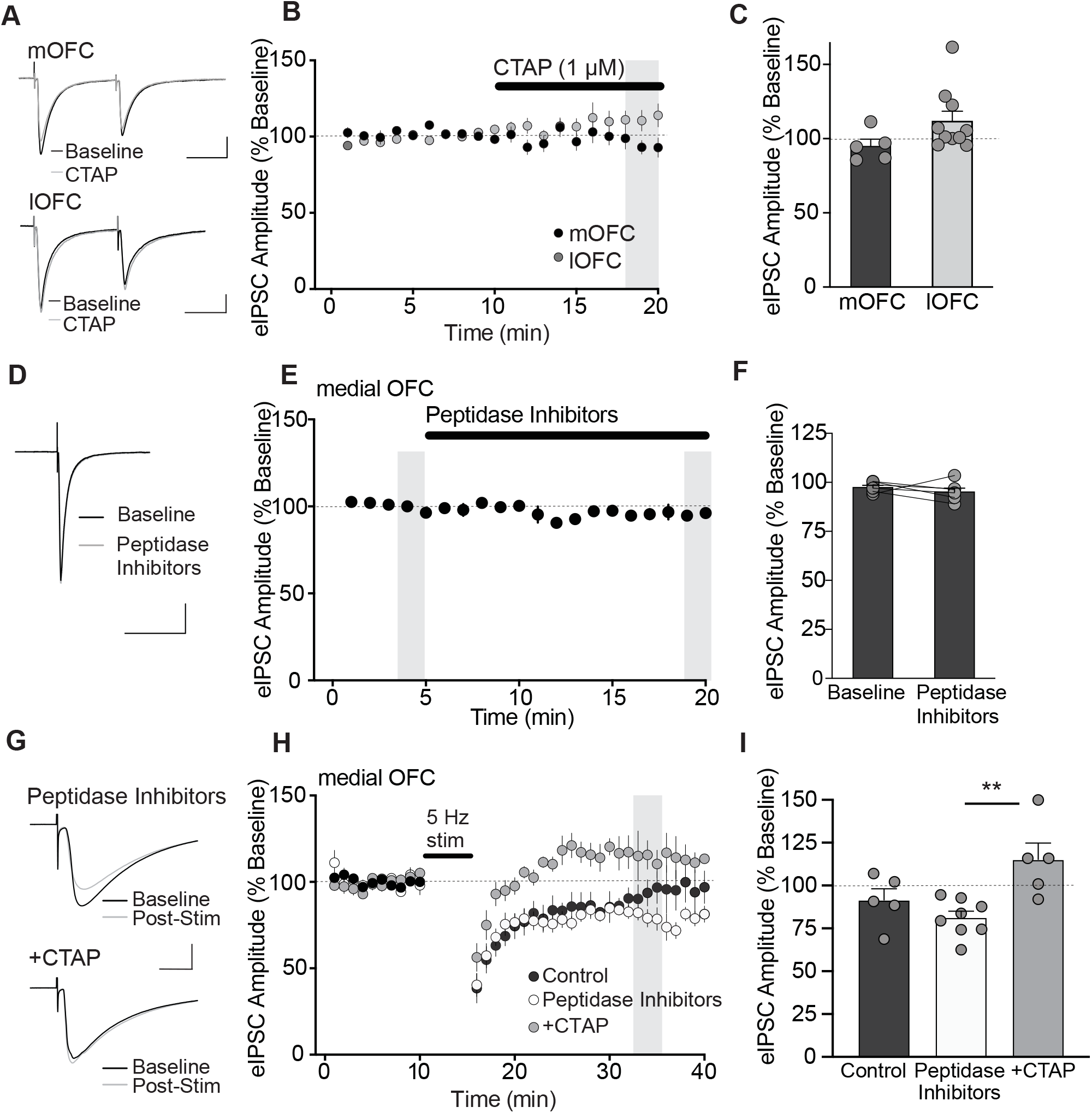
Endogenous opioids induced via physiological stimulation can induce long-term depression of GABA transmission. **A.** Averaged example traces of eIPSCs before (Baseline) and during CTAP (1 μM) application in mOFC or lOFC neurons. Scale bars: 250 pA, 25 ms (mOFC); 100 pA, 25 ms (lOFC). **B.** Time course of eIPSC amplitude before and during application of CTAP (1 μM) in mOFC or lOFC neurons. Application of CTAP did not alter eIPSC amplitude in either mOFC or lOFC pyramidal neurons. Shaded bar represents the sampled time point during CTAP application, which is quantified in C. **C**. Bar graph quantifying the change in eIPSC amplitude during CTAP application in mOFC and lOFC neurons. **D.** Averaged example traces of eIPSCs before (Baseline) and during application of the peptidase inhibitors, bestatin (10 μM), thiorphan (10 μM) and captopril (1 μM) in mOFC neurons. Scale bars: 250 pA, 25 ms. **E.** Time course of eIPSC amplitude before and during application of peptidase inhibitors in mOFC neurons. Shaded bars represent sampled time points before and during peptidase inhibitor application, which is quantified in C Blockade of peptide degradation did not affect basal eIPSC amplitude. **F.** Evoked IPSC amplitude before and during application of peptidase inhibitors. **G.** Averaged example traces illustrating eIPSCs before (Baseline) and after 5 Hz (5 min) synaptic stimulation in slices pre-treated with peptidase inhibitors alone or peptidase inhibitors with CTAP (1 μM). Scale bars, 500 pA, 10 ms. **H.** Time course of eIPSC amplitude before, during and after 5 Hz synaptic stimulation in mOFC neurons from control slices, and those pre-treated with peptidase inhibitors alone or peptidase inhibitors with CTAP (1 μM). Shaded bars represent sampled time points before and after 5 Hz stimulation, which are quantified in **I.** 5 Hz stimulation induced an irreversible, long-term suppression of eIPSC amplitude in peptidase inhibitor treated slices, but a reversible suppression in peptidase inhibitor + CTAP treated and control slices. I. Bar graph quantifying the percentage change in eIPSC amplitude following 5 Hz synaptic stimulation in mOFC neurons from control slices, or those pre-treated with peptidase inhibitors alone, or peptidase inhibitors with CTAP (1 μM). A Tukey’s posthoc test indicates a significant difference between responses in peptidase inhibitors and those with the addition of CTAP (*P<0.01).

Next, we examined whether endogenous opioid release induced via physiological stimulation could alter inhibitory synaptic transmission in the mOFC. In other brain regions, a moderate to high frequency stimulation can induce an opioid-induced LTD of excitatory or inhibitory synaptic transmission (Piskorowski and Chevaleyre, 2013; Wamsteeker Cusulin et al., 2013; Atwood et al., 2014). LTD at inhibitory synapses in the medial prefrontal cortex (mPFC) can be induced with moderate 5-10 Hz stimulation over 5 min in combination with neuromodulator receptor activation (Chiu et al., 2010). Given that exogenous opioid exposure can facilitate synaptic plasticity in our experiments, we hypothesized that enhancing endogenous opioid levels by preventing their metabolism could facilitate the induction of stimulation-induced inhibitory LTD. We first tested whether a cocktail of peptidase inhibitors alone could suppress inhibitory synaptic transmission. Application of bestatin (10 μM), thiorphan (10 μM) and captopril (1 μM) had no effect on eIPSC amplitude in the mOFC (t(5) = 0.09, P = 0.402; Fig. 8D-F). We next examined the effect of moderate frequency stimulation in the combined presence of peptidase inhibitors. A 5 Hz stimulation (5 min) induced an LTD of eIPSC amplitude lasting for at least 30 min (77 ± 5%, n/N = 8/6; Fig. 8G-I). This inhibitory LTD was blocked in slices pre-treated with CTAP (114 ± 8%, n/N = 5/3; Fig. 8G-I). Furthermore, in control slices without peptidase inhibitor treatment, 5 Hz stimulation did not induce a significant long-term suppression of evoked IPSC amplitude (96 ± 9%, n/N = 5/ 3, F(2,15) = 6.64, P = 0.0086; Fig. 8H-I). A Tukey’s multiple comparison test indicates a significant difference in responses in the presence of peptidase inhibitors and those with the addition of CTAP (P = 0.016, Fig. 8J). Together, these data indicate that under conditions of reduced metabolic degradation, stimulation of endogenous opioids can induce an LTD of inhibitory transmission onto mOFC neurons.

## Discussion

We found that mu-opioid receptor activation produces a long-lasting inhibition of GABAergic synaptic transmission onto layer II/III pyramidal neurons of the mOFC. This effect was regionally selective, as DAMGO did not alter inhibitory synaptic transmission onto layer II/III lOFC pyramidal neurons. Furthermore, inhibitory synaptic transmission from local PV+ inputs onto mOFC pyramidal neurons was sensitive to opioid modulation, indicating that the effect of mu-opioids are intrinsic to the OFC. Finally, endogenous opioids released by moderate frequency stimulation in the presence of protease inhibitors induced an LTD of inhibitory synaptic transmission onto mOFC pyramidal neurons.

### Subregional selectivity of mu-opioid action in the OFC

Although endogenous opioid expression was observed in both the lOFC and the mOFC, we observed DAMGO-induced suppression of inhibitory synaptic transmission only in layer II/III pyramidal neurons of the mOFC. A previous study indicated that DAMGO inhibits GABA release in the vlOFC of rats (Qu et al., 2015), a region that lies above the peak inflection of the rhinal sulcus (Van De Werd and Uylings, 2014). While we based our OFC subregions on the mouse atlas by Paxinos & Franklin (2001), in general our lOFC recordings were lateral to the vlOFC. Thus, there may be a medial-lateral gradient of mu-opioid sensitivity in the OFC. Future studies should examine if the projection targets of these subregions differ based on mu-opioid sensitivity. While we did not observe inhibition of GABA_A_ IPSCs onto layer II/III lOFC pyramidal neurons, it is possible that mu-opioids may influence inhibitory synaptic transmission in deeper layers (layer V) of the lOFC.

Our finding of subregional selectivity of mu-opioid action in the OFC is consistent with previous behavioural studies describing differential opioid-induced affective orofacial responses. Opioid hedonic hotspots are present in anterior regions of the mOFC and vOFC, whereas a cold spot is located in the posterior lOFC (Castro and Berridge, 2017). While the vOFC was not specifically examined in this study, the hotspot is localized more in the anterior and medial aspect of the OFC, whereas the coldspot is located more in the posterior and lateral aspect of the OFC, supporting differential opioid action between the mOFC and lOFC. Future work should examine if subregional differences in opioid modulation of inhibitory synaptic transmission may contribute to subregional differences in OFC-mediated behaviours (Izquierdo, 2017).

### Mu-opioids induce long-term depression of inhibitory synapses in the mOFC

To date, exogenous and endogenous opioid-induced LTD has only been demonstrated in a small number of studies (Iremonger and Bains, 2009; Chiu et al., 2010; Piskorowski and Chevaleyre, 2013; Wamsteeker Cusulin et al., 2013; Atwood et al., 2014). Consistent with these reports, we found that DAMGO induced a long-lasting suppression of inhibitory transmission. While pre-application of CTAP blocked the effect of DAMGO, application of CTAP after DAMGO-induced suppression of inhibition was ineffective, indicating that mechanisms engaged after receptor activation maintains the inhibition. Thus, DAMGO likely induced an LTD of inhibitory synapses onto mOFC pyramidal neurons. Several lines of evidence suggest that mu-opioid-induced LTD was mediated presynaptically. First, we observed a decrease in the frequency, but not amplitude of quantal mIPSCs, suggesting a decrease in the probability of release from GABA terminals. However, we observed no significant changes in paired-pulse ratio of evoked IPSCs. While this measure is typically associated with altered release probability, it can also measure postsynaptic diffusion of receptors (Blitz et al., 2004). A change in frequency of mIPSCs, but no alteration in paired-pulse ratio could suggest presynaptic silencing of GABA synapses by DAMGO; however this possibility requires examination using electron microscopy in future studies. Secondly, intracellular application of the PKA activator to the postsynaptic recording site did not influence DAMGO-induced LTD, yet extracellular application of forskolin impaired this effect, suggesting a presynaptic site of action. Finally, DAMGO-induced inhibition of electrically evoked IPSCs was similar to that when optically stimulating PV+ terminals, suggesting that the site of mu-opioid receptor activation is at PV+ inputs to mOFC pyramidal neurons.

### Mu opioid inhibitory long-term depression requires cAMP/PKA signalling

Previous work in the hippocampus (Lupica, 1995) or periaqueductal grey area (Vaughan and Christie, 1997) has indicated that mu-opioids suppress GABA release independent of presynaptic Ca^2+^ entry or G-protein coupled K^+^ channels, and may mediate its effect through direct interaction with the pre-synaptic release machinery at GABAergic terminals.

A direct downstream target of mu-opioid receptors is adenylyl cyclase, which is inhibited upon receptor activation, resulting in subsequent reduction of cAMP and PKA activity. Both cAMP/PKA have been implicated in multiple forms of presynaptic long-term plasticity, which are typically mediated by the endocannabinoid system (Chevaleyre et al., 2007; Mato et al., 2008; Tsetsenis et al., 2011)(Yang and Calakos, 2013). PKA is required in phosphorylating a number of presynaptic proteins involved in the induction and maintenance of LTD at inhibitory synapses, including the synaptic vesicle protein, Rab3B and the effector protein it binds, RIM1α (Chevaleyre et al., 2007; Tsetsenis et al., 2011). Given that both mu-opioid and CB1 receptors drive the Gα_i/o_ effector subunit of the G-protein signalling cascade, we surmised that mu-opioid LTD may also require cAMP and PKA. Consistent with this notion, we observed that enhancing cAMP/PKA activation with forskolin abolished mu-opioid LTD at inhibitory synapses onto mOFC pyramidal neurons. While our finding is consistent with previous reports showing that endocannabinoid-mediated forms of LTD depend on cAMP/PKA signalling (Chevaleyre et al., 2007; Mato et al., 2008; Tsetsenis et al., 2011), this is the first to demonstrate its requirement in LTD elicited by mu-opioid receptor activation. Furthermore, while forskolin application alone produced an expected facilitation of inhibitory transmission in mOFC pyramidal neurons, it had a negligible effect in lOFC pyramidal neurons, indicating varying sensitivity to cAMP/PKA. The lack of cAMP/PKA modulation in the lOFC did not appear to be due to occlusion of mu-opioid receptors, as we observed no functional endogenous opioid tone on these receptors. Together, our results suggest differential coupling of mu-opioid receptors to their downstream intracellular cascades in inhibitory synapses of the medial versus lateral OFC, likely explaining the varying effect of DAMGO in these subregions. While we have identified cAMP/PKA as a signalling pathway required for mu-opioid LTD, the downstream proteins involved in maintaining this plasticity have yet to be determined. Further work using genetic strategies would be needed to investigate the role of Rab3B and RIM1α.

### Mu-opioids modulate synaptic transmission from local PV+ inputs to mOFC neurons

In the hippocampus, perisomatically projecting PV-expressing basket cells exhibit a much higher percentage of colocalization with mu-opioid receptors than other interneurons (Drake and Milner, 2002, 2006). In contrast, mu-opioid receptor immunoreactivity was primarily colocalized with vasoactive intestinal polypeptide (VIP)-containing neurons and to a lesser extent on PV+ interneurons in layers ll-IV of the frontopariatal cortex (Taki et al., 2000). We found that mu-opioids inhibit GABAergic synaptic transmission from PV+ inputs onto mOFC pyramidal neurons. This is likely due to a direct effect of mu-opioid stimulation at PV+ terminals. These experiments were performed in the presence of DNQX to block excitatory synaptic transmission, thereby excluding the possibility of a heterosynaptically-mediated suppression of inhibitory inputs to pyramidal neurons. Given that the maximal inhibition of oIPSCs from local PV+ inputs by DAMGO was similar to that produced by bulk electrical stimulation of non-specific local and external GABA inputs, we propose that the majority of mu-opioid suppression of inhibitory transmission is mediated by local PV+ terminals, suggesting that mu-opioids can act intrinsically within the mOFC. Consistent with our findings, perisomatically projecting basket cells in the hippocampus are twice as likely to be hyperpolarized by mu-opioid receptors than dendritically projecting interneurons (Svoboda et al., 1999). Because PV+ interneurons are required for generation of gamma oscillations in the prefrontal cortex (Sohal et al., 2009), changes in mu-opioid action may play a significant role in modulating cortical network performance. PV+ interneurons in the OFC are critical for cognitive flexibility (Bissonette et al., 2015). Furthermore, inhibition of mu-opioid receptors in the OFC of male mice impaired behavioural flexibility on a spatial reversal learning task (Laredo et al., 2015). Thus, it is feasible that mu-opioid modulation of PV+ interneuron-mediated inhibitory synaptic transmission onto mOFC pyramidal neurons may contribute to optimized behavioural flexibility.

### Endogenous opioid stimulation elicits long-term depression of inhibitory synapses in the mOFC

Here, we have demonstrated that moderate frequency stimulation in the mOFC can induce an LTD of inhibitory transmission dependent on endogenous opioid release and activation of mu-opioid receptors. This suppression of inhibition was transient under naive conditions, but became persistent in the presence of peptidase inhibitors. This indicates that under normal conditions, moderate to high levels of activity can induce endogenous opioid release, but metabolic degradative enzymes, at least in-vitro, tightly restrict their action. It is important to note that the targeted peptidases are non-selective metalloproteases and may have the potential to enhance the activity of other non-opioid endogenous signalling peptides, for example, substance P and neurokinin B (Iritani et al., 1989; Wang et al., 1991; Marksteiner et al., 1992), that may act either in concert with or opposition to endogenous opioid signalling. While we did not observe a significant change in evoked IPSCs with peptidase inhibitors alone, we cannot exclude that other endogenous peptides may be modulating opioid-inhibition of IPSCs. It remains unclear which endogenous opioids mediate mu opioid-LTD at inhibitory synapses, as both β-endorphin and met-enkephalin are expressed in the mOFC. Notably, β-endorphin is much less sensitive to peptidase degradation compared to met-enkephalin (McKnight et al., 1982). Furthermore, while we determined that β-endorphin is not co-localized with PV+ interneurons, further work could determine which subpopulation of cells in the OFC express these endogenous opioids. In summary, we have demonstrated that endogenous opioid release can be induced by physiological stimulation, but only elicits a functional effect on inhibitory synapses in the mOFC dependent on metabolic activity.

Our work is consistent with previous evidence demonstrating endogenous opioid-induced LTD at inhibitory synapses. Within the hypothalamus, kappa opioid-mediated LTD depends on retrograde opioid signalling, induced via postsynaptic depolarization and activation of mGluR5 (Wamsteeker Cusulin et al., 2013). While the locus of opioid release was not determined in the present study, retrograde opioid signalling was unlikely given that mOFC pyramidal neurons were maintained at −70 mV during the stimulation protocol. In the striatum, mu-opioid induced LTD at inhibitory synapses also occurs presynaptically and in the presence of a peptidase inhibitor (Atwood et al., 2014). Similarly, a presynaptically-mediated delta opioid-LTD at inhibitory synapses in the CA2 of the hippocampus can occur (Piskorowski and Chevaleyre, 2013). Thus, opioid-induced LTD can involve different opioid release mechanisms and/or variations in the downstream intracellular cascades activated by mu, kappa and delta opioid receptor activation in different brain regions.

### Conclusions

Our findings elucidate the cellular mechanism of action by mu-opioids in the OFC. We found subregional selectivity of mu-opioid action such that mu-opioids supressed inhibition in the mOFC rather than the lOFC. This sub-regional difference may be associated with the differential functional effects by mu-opioids on OFC dependent behaviours. Mu-opioid suppression of inhibitory synaptic transmission in the mOFC was long lasting, suggesting a role of mu-opioids in mediating long-term adaptive changes within this region. Furthermore, these mu-opioid effects were in large part mediated by PV+ interneurons in the OFC, and therefore, may play a role in shaping cortical oscillatory activity. Given that activity-dependent endogenous opioid release can suppress inhibitory synaptic transmission in the OFC, this effect may play a fundamental role in shifting network dynamics during reward-seeking behaviours. While these studies primarily focused on OFC from male mice, future studies could examine if similar mechanisms of opioid action occur in the OFC of female mice. In conclusion, understanding how opioids influence synaptic transmission within the OFC may reveal new mechanisms of action and potential treatments for multiple psychiatric disorders, where altered mu-opioid signaling within the OFC may lead to impaired behavioral flexibility and a loss of inhibitory control over appetitively motivated behaviors such as observed in drug abuse and binge eating disorders.

## Acknowledgements

This work is supported by a Tier 1 Canada Research Chair and Canada Institutes for Health Research Foundation Grant (CIHR FDN 148473 SLB). BA was supported by a Cumming School of Medicine Graduate Scholarship. BKL was supported by an Alberta Innovates Health Solutions Postdoctoral Fellowship. The authors wish to acknowledge the advanced microscopy centre at the Hotchkiss Brain Institute for use of their confocal and cryostat.

## Author Contributions

BA and BKL performed electrophysiological experiments. MQ and BA performed immunohistochemistry experiments. CST performed intracranial virus injections. BA, BKL and SLB wrote first drafts of the manuscript. BKL, BA, CST and SLB revised the manuscript.

---

**Extended Data Figure 7-1.** Representative images of met-enkaphalin-like immunofluorescence in the mOFC, lOFC and striatum. Scale bar 100 μm.

## References

Atwood BK, Kupferschmidt DA, Lovinger DM (2014) Opioids induce dissociable forms of longterm depression of excitatory inputs to the dorsal striatum. Nat Neurosci 17:540–548.

Badiani A, Leone P, Noel MB, Stewart J (1995) Ventral tegmental area opioid mechanisms and modulation of ingestive behavior. Brain Res 670:264–276.

Bissonette GB, Schoenbaum G, Roesch MR, Powell EM (2015) Interneurons are necessary for coordinated activity during reversal learning in orbitofrontal cortex. Biol Psychiatry 77:454–464.

Blitz DM, Foster KA, Regehr WG (2004) Short-term synaptic plasticity: a comparison of two synapses. Nat Rev Neurosci 5:630–640.

Castro DC, Berridge KC (2017) Opioid and orexin hedonic hotspots in rat orbitofrontal cortex and insula. Proc Natl Acad Sci:201705753.

Chevaleyre V, Heifets BD, Kaeser PS, Südhof TC, Castillo PE (2007) Endocannabinoid-Mediated Long-Term Plasticity Requires cAMP/PKA Signaling and RIM1α. Neuron 54:801–812.

Chiu CQ, Puente N, Grandes P, Castillo PE (2010) Dopaminergic modulation of endocannabinoid-mediated plasticity at GABAergic synapses in the prefrontal cortex. J Neurosci Off J Soc Neurosci 30:7236–7248.

Drake CT, Milner TA (2002) Mu opioid receptors are in discrete hippocampal interneuron subpopulations. Hippocampus 12:119–136.

Drake CT, Milner TA (2006) Mu opioid receptors are extensively co-localized with parvalbumin, but not somatostatin, in the dentate gyrus. Neurosci Lett 403:176–180.

Gourley SL, Lee AS, Howell JL, Pittenger C, Taylor JR (2010) Dissociable regulation of instrumental action within mouse prefrontal cortex. Eur J Neurosci 32:1726–1734.

Gremel CM, Costa RM (2013) Orbitofrontal and striatal circuits dynamically encode the shift between goal-directed and habitual actions. Nat Commun 4:2264.

Hoover WB, Vertes RP (2011) Projections of the medial orbital and ventral orbital cortex in the rat. J Comp Neurol 519:3766–3801.

Hunnicutt BJ, Long BR, Kusefoglu D, Gertz KJ, Zhong H, Mao T (2014) A comprehensive thalamocortical projection map at the mesoscopic level. Nat Neurosci 17:1276–1285.

Iremonger KJ, Bains JS (2009) Retrograde opioid signaling regulates glutamatergic transmission in the hypothalamus. J Neurosci Off J Soc Neurosci 29:7349–7358.

Iritani S, Fujii M, Satoh K (1989) The distribution of substance P in the cerebral cortex and hippocampal formation: an immunohistochemical study in the monkey and rat. Brain Res Bull 22:295–303.

Izquierdo A (2017) Functional Heterogeneity within Rat Orbitofrontal Cortex in Reward Learning and Decision Making. J Neurosci Off J Soc Neurosci 37:10529–10540.

Krettek JE, Price JL (1977) The cortical projections of the mediodorsal nucleus and adjacent thalamic nuclei in the rat. J Comp Neurol 171:157–191.

Laredo SA, Steinman MQ, Robles CF, Ferrer E, Ragen BJ, Trainor BC (2015) Effects of defeat stress on behavioral flexibility in males and females: modulation by the mu-opioid receptor. Eur J Neurosci 41:434–441.

Little JP, Carter AG (2012) Subcellular synaptic connectivity of layer 2 pyramidal neurons in the medial prefrontal cortex. J Neurosci Off J Soc Neurosci 32:12808–12819.

Lupica CR (1995) Delta and mu enkephalins inhibit spontaneous GABA-mediated IPSCs via a cyclic AMP-independent mechanism in the rat hippocampus. J Neurosci Off J Soc Neurosci 15:737–749.

MacDonald AF, Billington CJ, Levine AS (2004) Alterations in food intake by opioid and dopamine signaling pathways between the ventral tegmental area and the shell of the nucleus accumbens. Brain Res 1018:78–85.

Madison DV, Nicoll RA (1988) Enkephalin hyperpolarizes interneurones in the rat hippocampus. J Physiol 398:123–130.

Malvaez M, Shieh C, Murphy MD, Greenfield VY, Wassum KM (2019) Distinct cortical-amygdala projections drive reward value encoding and retrieval. Nat Neurosci 22:762–769.

Mansour A, Fox CA, Burke S, Akil H, Watson SJ (1995) Immunohistochemical localization of the cloned mu opioid receptor in the rat CNS. J Chem Neuroanat 8:283–305.

Marksteiner J, Sperk G, Krause JE (1992) Distribution of neurons expressing neurokinin B in the rat brain: immunohistochemistry and in situ hybridization. J Comp Neurol 317:341–356.

Mato S, Lafourcade M, Robbe D, Bakiri Y, Manzoni OJ (2008) Role of the cyclic-AMP/PKA cascade and of P/Q-type Ca++ channels in endocannabinoid-mediated long-term depression in the nucleus accumbens. Neuropharmacology 54:87–94.

McKnight AT, Corbett AD, Paterson SJ, Magnan J, Kosterlitz HW (1982) Comparison of in vitro potencies in pharmacological and binding assays after inhibition of peptidases reveals that dynorphin (1-9) is a potent kappa-agonist. Life Sci 31:1725–1728.

McQuiston AR, Saggau P (2003) Mu-opioid receptors facilitate the propagation of excitatory activity in rat hippocampal area CA1 by disinhibition of all anatomical layers. J Neurophysiol 90:1936–1948.

Mena JD, Sadeghian K, Baldo BA (2011) Induction of hyperphagia and carbohydrate intake by μ-opioid receptor stimulation in circumscribed regions of frontal cortex. J Neurosci Off J Soc Neurosci 31:3249–3260.

Mucha RF, Iversen SD (1986) Increased food intake after opioid microinjections into nucleus accumbens and ventral tegmental area of rat. Brain Res 397:214–224.

Murphy MJM, Deutch AY (2018) Organization of afferents to the orbitofrontal cortex in the rat. J Comp Neurol 526:1498–1526.

Namboodiri VMK, Otis JM, Heeswijk K van, Voets ES, Alghorazi RA, Rodriguez-Romaguera J, Mihalas S, Stuber GD (2019) Single-cell activity tracking reveals that orbitofrontal neurons acquire and maintain a long-term memory to guide behavioral adaptation. Nat Neurosci 22:1110.

Nummenmaa L, Saanijoki T, Tuominen L, Hirvonen J, Tuulari JJ, Nuutila P, Kalliokoski K (2018) μ-opioid receptor system mediates reward processing in humans. Nat Commun 9:1500.

Ongür D, Price JL (2000) The organization of networks within the orbital and medial prefrontal cortex of rats, monkeys and humans. Cereb Cortex N Y N 1991 10:206–219.

Paxinos G, Franklin KBJ (2001) The mouse brain in stereotaxic coordinates. 2nd Edition, Academic Press, San Diego.

Piskorowski RA, Chevaleyre V (2013) Delta-Opioid Receptors Mediate Unique Plasticity onto Parvalbumin-Expressing Interneurons in Area CA2 of the Hippocampus. J Neurosci 33:14567–14578.

Qu C-L, Huo F-Q, Huang F-S, Tang J-S (2015) Activation of mu-opioid receptors in the ventrolateral orbital cortex inhibits the GABAergic miniature inhibitory postsynaptic currents in rats. Neurosci Lett 592:64–69.

Quirk MC, Sosulski DL, Feierstein CE, Uchida N, Mainen ZF (2009) A defined network of fastspiking interneurons in orbitofrontal cortex: responses to behavioral contingencies and ketamine administration. Front Syst Neurosci 3:13.

Ray JP, Price JL (1992) The organization of the thalamocortical connections of the mediodorsal thalamic nucleus in the rat, related to the ventral forebrain-prefrontal cortex topography. J Comp Neurol 323:167–197.

Sohal VS, Zhang F, Yizhar O, Deisseroth K (2009) Parvalbumin neurons and gamma rhythms enhance cortical circuit performance. Nature 459:698–702.

Svoboda KR, Adams CE, Lupica CR (1999) Opioid receptor subtype expression defines morphologically distinct classes of hippocampal interneurons. J Neurosci Off J Soc Neurosci 19:85–95.

Taki K, Kaneko T, Mizuno N (2000) A group of cortical interneurons expressing mu-opioid receptor-like immunoreactivity: a double immunofluorescence study in the rat cerebral cortex. Neuroscience 98:221–231.

Tsetsenis T, Younts TJ, Chiu CQ, Kaeser PS, Castillo PE, Südhof TC (2011) Rab3B protein is required for long-term depression of hippocampal inhibitory synapses and for normal reversal learning. Proc Natl Acad Sci U S A 108:14300–14305.

Van De Werd HJJM, Uylings HBM (2014) Comparison of (stereotactic) parcellations in mouse prefrontal cortex. Brain Struct Funct 219:433–459.

Vaughan CW, Christie MJ (1997) Presynaptic inhibitory action of opioids on synaptic transmission in the rat periaqueductal grey in vitro. J Physiol 498 (Pt 2):463–472.

Wamsteeker Cusulin JI, Füzesi T, Inoue W, Bains JS (2013) Glucocorticoid feedback uncovers retrograde opioid signaling at hypothalamic synapses. Nat Neurosci 16:596–604.

Wang LH, Ahmad S, Benter IF, Chow A, Mizutani S, Ward PE (1991) Differential processing of substance P and neurokinin A by plasma dipeptidyl(amino)peptidase IV, aminopeptidase M and angiotensin converting enzyme. Peptides 12:1357–1364.

Yang Y, Calakos N (2013) Presynaptic long-term plasticity. Front Synaptic Neurosci 5 Available at: https://www.ncbi.nlm.nih.gov/pmc/articles/PMC3797957/ [Accessed April 18, 2020].

Zieglgänsberger W, French ED, Siggins GR, Bloom FE (1979) Opioid peptides may excite hippocampal pyramidal neurons by inhibiting adjacent inhibitory interneurons. Science 205:415–417.

