## Supplementary figures and images for "Mu-opioids suppress GABAergic synaptic transmission onto orbitofrontal cortex pyramidal neurons with subregional selectivity"

### Figure S1

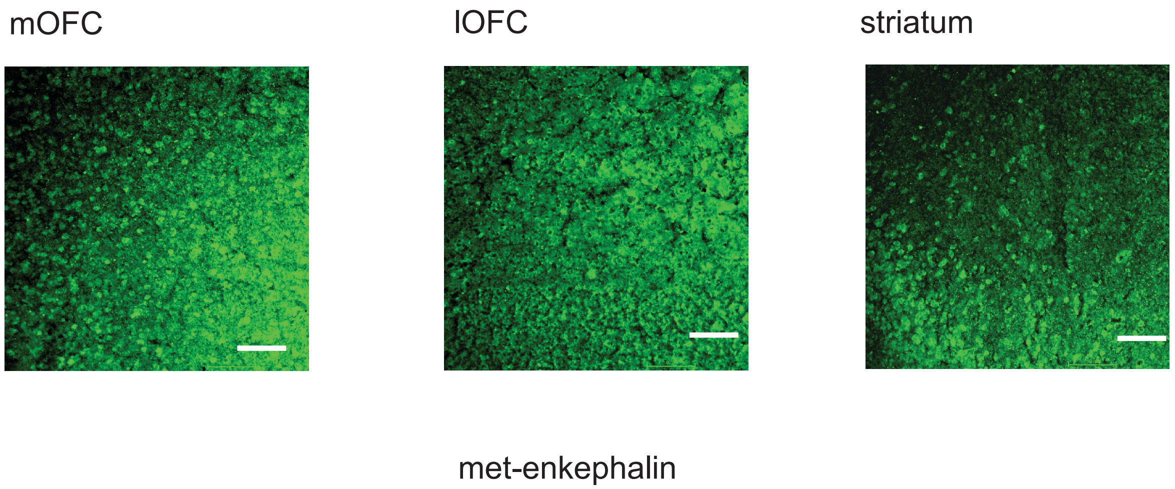
